# Dissection of barrier dysfunction in organoid-derived human intestinal epithelia induced by *Giardia duodenalis*

**DOI:** 10.1101/2020.11.17.384537

**Authors:** Martin Kraft, David Holthaus, Susanne M. Krug, Gudrun Holland, Joerg-Dieter Schulzke, Toni Aebischer, Christian Klotz

**Affiliations:** Department of Infectious Diseases, Unit 16 Mycotic and Parasitic Agents and Mycobacteria, Robert Koch-Institute, Berlin, Germany; Department of Gastroenterology, Rheumatology and Infectious Diseases, Institute of Clinical Physiology/Nutritional Medicine, Charité - Universitätsmedizin Berlin, Berlin, Germany; Advanced Light and Electron Microscopy, Centre for Biological Threats and Special Pathogens, Robert Koch Institute

**Keywords:** Giardiasis, host-parasite interaction, pathogenesis of intestinal protozoa

## Abstract

**Background and aims:** The protozoa *Giardia duodenalis* is a major cause of gastrointestinal illness worldwide, but underlying pathophysiological mechanisms remain obscure, partly due to the absence of adequate cellular models. We aimed to overcome these limitations and to recapitulate the authentic series of events in the primary human duodenal tissue by using the human organoid system.

**Methods:** We established a compartmentalized cellular transwell system with electrophysiological and barrier properties akin to duodenal mucosa and dissected the events leading to *G. duodenalis*-induced barrier breakdown by functional analysis of transcriptional, electrophysiological and tight junction components.

**Results:** Organoid-derived cell layers of different donors showed a time- and parasite load-dependent leak flux indicated by collapse of epithelial barrier upon *G. duodenalis* infection. Transcriptomic analysis suggested major expression changes in genes contributing to ion transport and tight junction structure. SLC12A2/NKCC1- and CFTR-dependent chloride secretion was reduced early after infection, while changes in the tight junction composition, localization and structural organization occurred later as revealed by immunofluorescence analysis and freeze fracture electron microscopy.

**Conclusion:** Data suggest a previously unknown sequence of events culminating in intestinal barrier dysfunction upon *G. duodenalis* infection ignited by alterations of cellular ion transport followed by breakdown of the tight junctional complex and loss of epithelial integrity. The newly established organoid-derived model to study *G. duodenalis* infection will help enable further molecular dissection of the disease mechanism and, thus, can help to find new options treating disease and infection, in particular relevant for chronic cases of giardiasis.

## Introduction

Protozoan parasites of the species complex *Giardia duodenalis* cause the gastrointestinal disease giardiasis.^1^ It is one of the most prevalent parasitic diseases worldwide with an estimated number of about 183 million symptomatic cases each year.^2^ The parasites are non-invasive and colonize the small intestine, where the trophozoites, the disease causing form, replicates. During the intestinal passage the parasites transform into the environmentally resistant cyst stage, which is transmitted to the next host via the fecal oral route. Disease mechanisms in humans are not understood, partly due to the lack of adequate model systems.

The clinical outcome which varies between individuals, ranging from asymptomatic colonization to severe acute or chronic disease, is thought to reflect a complex interaction of *Giardia* parasites with the host epithelium, modulated by the local microbiome and dietary factors.^1, 3, 4^ Common symptoms are malabsorption, diarrhea, bloating, abdominal pain, nausea and other gastrointestinal complaints.^1^ Long term sequelae that have been linked to infection are stunting (in pediatric patients), irritable bowel disease, and chronic fatigue syndrome.^5, 6^

There are only few studies reporting on histological correlates of infection in asymptomatic and symptomatic, acute or chronic disease. In a large case series of 567 adult patients undergoing endoscopy because of unspecific gastrointestinal complaints, giardiasis was diagnosed in biopsies of the gastrointestinal tract where parasites were found primarily in the duodenum. There, signs of an inflammatory reaction were detected in less than 4% of cases and the mucosa seemed unaffected in the vast majority of patients.^7^ In contrast, another to date unmatched study on 13 patients suffering from chronic symptomatic giardiasis showed increased recruitment of inflammatory cells to the mucosa, a 50% reduction of villous surface area and decreased electrical resistance of biopsies, indicative of compromised absorptive and barrier properties of the epithelium.^8^

To date, attempts to reproduce these *in vivo* findings using common *in vitro* models of intestinal epithelia to decipher the molecular mechanisms behind and identify responsible parasite, host or other factors, respectively, have yielded conflicting results.^1, 9^ More recently, stem cell derived primary organoid culture models have been developed and represent a promising tool to improve the *in vitro* model systems to study host *Giardia* interaction.^10–12^

In this study, we established a reliable *in vitro* system based on human stem cell-derived organoids to decipher the effects of *G. duodenalis* infection on barrier function in human primary tissue under conditions that preserve *G. duodenalis* growth potential while overcoming their negative effect observed in host cell survival when using conventional cell models. This enabled the elucidation of a parasite-induced cascade of events: Transcriptomic, functional and structural analysis revealed an early decrease of ion transporter mRNA abundance leading to reduced channel activity, followed by epithelial barrier breakdown due to modulation of tight junction composition, localization and structural organization.

## Methods

A further description of material and methods is available in supplementary methods.

### Organoid culture

Human organoids were generated and maintained from duodenal biopsies from healthy volunteers undergoing routine exams at Charité, University Hospital, Berlin (approved by local authorities, #EA4-015-13) as previously described (details in Supplementary materials).^13–16^ Organoid-derived monolayers (ODMs) were prepared in Matrigel-coated transwell cell culture inserts (Merck Millipore, 0.6 cm^2^, 0.4 μm pores). For this, organoids were harvested in ice-cold AdvDMEM/F12, centrifuged and mechanically disrupted as described for passaging. Cells were sedimented, resuspended in differentiation medium (organoid medium without WRN-CM, A83-01 and SB202190) and added to the apical compartment of the cell insert (two 50 μl Matrigel droplets of 3D-culture were required for one transwell insert with an area of 0.6 cm^2^). Differentiation medium was applied to the upper and lower compartment of the transwell system and exchanged every 2-3 days. For the first two days 10 μM Y-27632 was added to inhibit anoikis.

### Parasite culture

*G. duodenalis* WB6 (ATCC 50803) trophozoites were cultured at 37 °C in flat-sided 10-ml-tubes (Nunclon, Thermo Scientific) in Keister’s modified TYI-S-33 medium^17^, supplemented with 10% adult bovine serum (Gibco), 100 μg/mL streptomycin/100 U/mL penicillin (Capricorn) and 0,05% bovine/ovine bile (Sigma). For passaging, culture tubes were put on ice for 20 min to facilitate trophozoite detachment. Trophozoits were passaged the day before infection experiments to guarantee logarithmic growth phase. For infection experiments cells were quantified using a Neubauer counting chamber (C-Chip, Biochrom AG) and pelleted at 1000 × *g* for 5 min at 4°C. The pellet was suspended according to desired amounts in complete TYI-S-33 medium.

### Infection of ODMs

The apical compartment of 8-10 days old ODMs was replaced with complete TYI-S-33 the evening before infection to ensure adaption of the cells to TYI-S-33 medium. Medium was renewed again before infection in both compartments, i.e. TYI-S-33 in the apical and organoid differentiation medium in the basal compartment. Finally, *G. duodenalis* trophozoites were added into the apical compartment. For lysate experiments, trophozoites were sonicated for 10 min at 4°C (30s ON/OFF) and a power of 72 D with a 450 Digital Sonifier (Branson) and subsequently added to the monolayers.

### Transepithelial electric resistance (TEER) measurements

TEER measurements, used for barrier dysfunction experiments were conducted on a 37 °C heating block using a Millicell ERS-2 Voltohmmeter (Merck-Millipore) equipped with an Ag/AgCl electrode (STX01). Blank electric resistance (cell-free transwell insert) was subtracted from raw resistance and standardized for 1 cm² surface area.

### Dilution potential measurements

Dilution potential measurements to determinate permeabilities for sodium and chloride were performed in Ussing chambers modified for cell-culture inserts. Water-jacketed gas lifts kept at 37 °C were filled with 10 mL bathing solution that contained (in mM) 119 NaCl, 21 NaHCO_3_, 5.4 KCl, 1.2 CaCl_2_, 1 MgSO_4_, 3 HEPES, and 10 D(+)-glucose, and was gassed with 95% O_2_ and 5% CO_2_ resulting in a pH of 7.4. Data were corrected for the resistance of the empty filter and the bathing solution. Dilution potentials were measured replacing half of the NaCl by mannitol (iso-osmotical solution) on the apical or basolateral side of the epithelial monolayer. The ratio of P_Na_ and P_Cl_ and the absolute permeabilities for Na^+^ and Cl^−^ were calculated using the Goldman-Hodgkin-Katz equation ^5^.

### Short-circuit current and ion secretion/transport via the NKCC and CFTR

Determination of net ion transport was done by measurement of changes in short-circuit current (I_SC_) until reaching a steady state. Secretion of Cl^−^ by monolayers was stimulated with PGE2 (1 μM, basal side) and theophylline (10 mM, both sides). The effect of theophylline and PGE2 was antagonized by bumetanide (10 μM, basal side) inhibiting the NKCC and ΔI_SC_ was received by comparison of both conditions. To determine the activity of the CFTR similar experiments were performed stimulating chloride secretion by 10 μM forskolin, inhibiting the CFTR with 10 μM 5-nitro-2-(3-phenylpropylamino) benzoic acid (NPPB) and stimulating again, resulting in only unspecific secretion.

### Fluorescein flux assay

Fluorescein (332 Da) was used to investigate permeability for a still small, but already bigger than main ions molecules using the Ussing-chamber setup under short-circuit conditions. For paracellular flux measurements, fluorescein (0.1 mM) was added apically and basal samples were taken at 0, 10, 20, 30, and 40 min. Tracer fluxes and apparent permeabilities were calculated from the amount of fluorescein in the basal compartment measured fluorometrically by Tecan Infinite® M200 plate reader (excitation 485 nm; emission 525 nm; bandwidth 20 nm).

### Quantitative real time PCR

RNA was extracted from organoid-derived monolayer (ODM) and organoid cultures using Direct-zol RNA Microprep kit (Zymo) followed by on-column DNase I treatment following manufacturer’s protocol. RNA was quantified at 260/280 nm using Infinite M200 Pro plate reader (Tecan). RNA was reversely transcribed using High Capacity RNA-to-cDNA Kit (Applied Biosystems) with 500 ng per reaction. RT-qPCR was performed with a C1000 cycler with CFX96 head (Biorad), and included an initial 10 min enzyme activation step at 95 °C, followed by 40 cycles of 20 s at 95 °C, 30 s at 60 °C and 20 s at 72 °C. Melting curve analysis was performed to verify amplicon specificity. Relative expression compared to control cells was calculated using the ΔΔCT method. Primer sequences are included in table S1 of the supplementary material.

## Results

### A human two-dimensional primary intestinal epithelium suitable to model *G. duodenalis* infections

To study the effects of *G. duodenalis* on barrier function of human primary duodenal epithelial cells we established a suitable two-dimensional model that fulfills the following criteria: 1) apical and basolateral compartments separated by a polarized columnar cell layer, enabling separate manipulation of the compartments; 2) electrophysiological and barrier properties akin to a duodenal epithelium.

Using three dimensional organoid cultures from human duodenal biopsies, we set-up a two-dimensional transwell system, referred to as organoid-derived monolayers (ODM). ^14, 16^ To induce differentiation in ODM, we omitted the stem cell promoting factors Wnt3a, A83 (TGF-β inhibitor) and SB202190 (p38 inhibitor) that are part of the stem cell enrichment medium (see methods for details). ODMs reproducibly showed increasing TEER values that reached a plateau of ~ 250 Ω*cm^2^ after 9 days of culture (Figure 1A). At this time point, cells showed a typical polarized enterocyte-like shape with microvilli and expression of transporter proteins at their apical tips (Figure 1B) and an approximate height of ~20 μm (Figure 1C). Transmission electron microscopy of ultra-thin sections revealed well-developed cellular junction complexes, including TJs, adherence junctions and desmosomes (Figure 1D). We further characterized the ODMs by determining relative abundance of cell-type specific marker gene transcripts in comparison to the initial stem-cell enriched organoid culture. This revealed enrichment for enterocyte/absorptive cells while secretory cell types (i.e., Paneth cells, goblet cells, enteroendocrine cells) were reduced (Figure 1E). Immunostaining to detect cell type specific marker proteins corroborated these findings (supplementary Figure 1). Altogether, this implies the establishment of a tight epithelial barrier composed by a primary enterocyte-type cell population, the dominant cell type of the natural duodenal epithelium.

**Figure 1:**
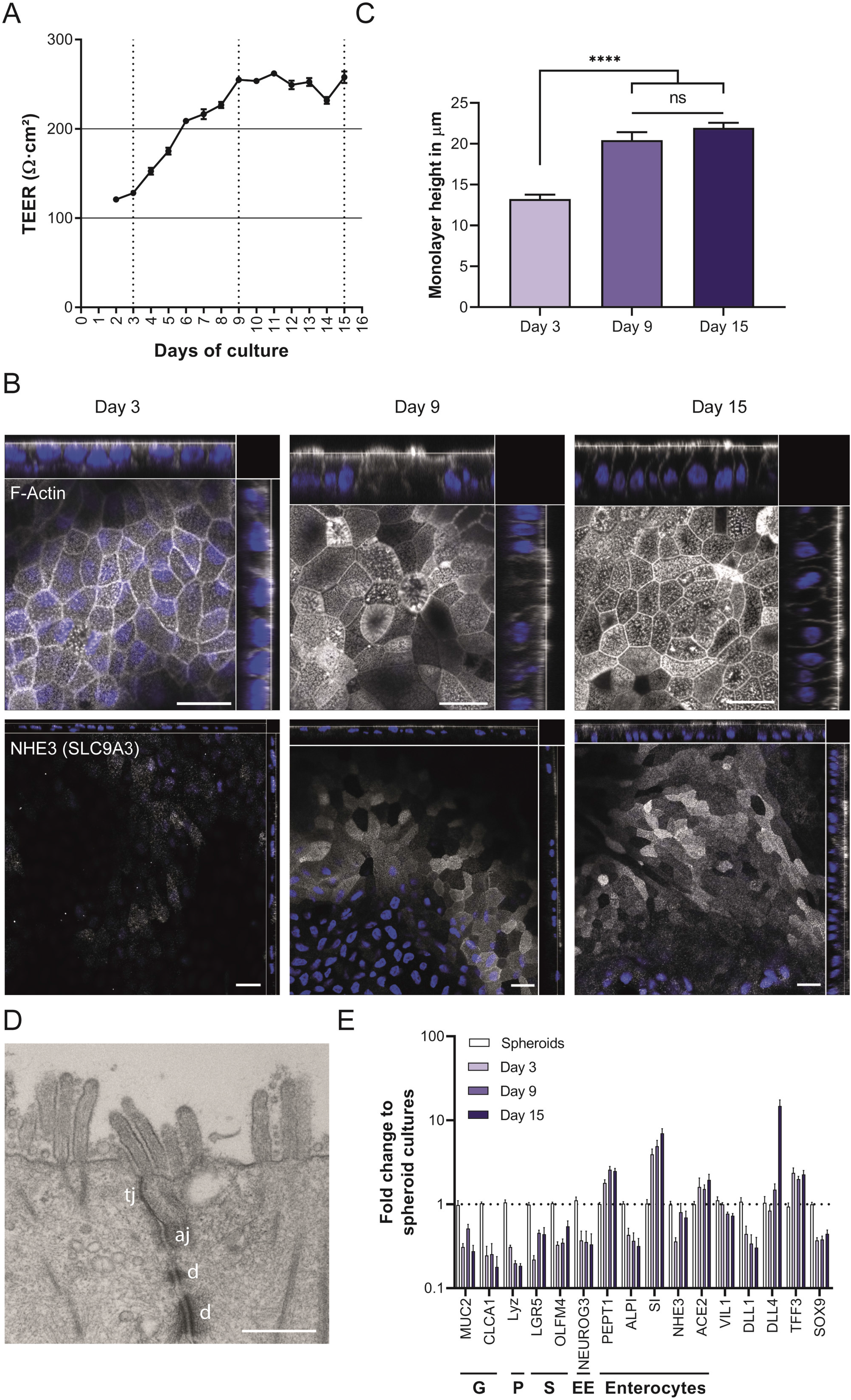
Characterization of human intestinal organoid-derived monolayers (ODMs). Human duodenal organoids were seeded on transwell filters to generate electrophysiologically tight epithelia, reaching (A) a plateau of transepithelial-electric resistance (TEER) 9 days after seeding of approximately 250 Ω.cm^2^. (B) Representative (immuno)-fluorescence microscopic analysis of ODMs on days 3, 9 and 15 after seeding show apical staining of brush border F-actin with Phalloidin and enterocyte-typical transporter NHE3 (SLC9A3). Cells become more columnar and express NHE3 localized at the apical brush border. Orthogonal stacks are shown. Scale bars represent 20 μm. (C) Thickness of ODMs significantly increases with culture time reaching a stable plateau 9 days after seeding and forming a columnar cell layer of approximately 20μm height. (D) Transmission electron microscopy of an ultrathin section through an ODM monolayer. The image shows the apical contact zone of two cells which reveal the presence of a tight junction (tj), followed by an adherens junction (aj) and desmosomes (d). Scale bars represent 500 nm. (E) Characterization of ODMS by RT-qPCR for various marker genes representing the intestinal cell types enteroendocrine cells (EE), Paneth cells (P), goblet cells (G), stem cells (S) and enterocytes on days 3, 9 and 15 after seeding. Expression levels were normalized to stem cell-enriched organoid cultures and showing higher abundance of enterocyte-typical gene expression. All quantitative experiments show mean (± SEM) of at least seven individual transwell filters of a minimum of three independent experiments. Statistical significance of the height difference of the cell layers of various time points was determined using ANOVA with Tuckey’s correction for multiple testing **** p < 0.0001

### Infection of human primary intestinal epithelium with G. duodenalis leads to barrier breakdown

For *in vitro* growth, *G. duodenalis* as a microaerophilic organism requires a highly reducing growth medium. Keisters’ modified TYI-S-33 is used for this purpose. In addition to tryptone, yeast extract and ferric ammonium citrate, it contains bile acids to better mimic duodenal conditions. This medium was found to be toxic to cell lines such as commonly used human colon carcinoma-derived Caco-2 cells and, therefore, respective studies often used medium mixtures or pure DMEM-based medium to perform co-culture experiments *in vitro*.^9, 18^ However, ODM exposed to TYI-S-33 in the upper, apical compartment of the transwell system and organoid differentiation medium in the basolateral compartment, showed stable TEER values after equilibration for at least 3 days (Figure 2A, 2B). This enabled the use of these conditions for subsequent experiments.

**Figure 2:**
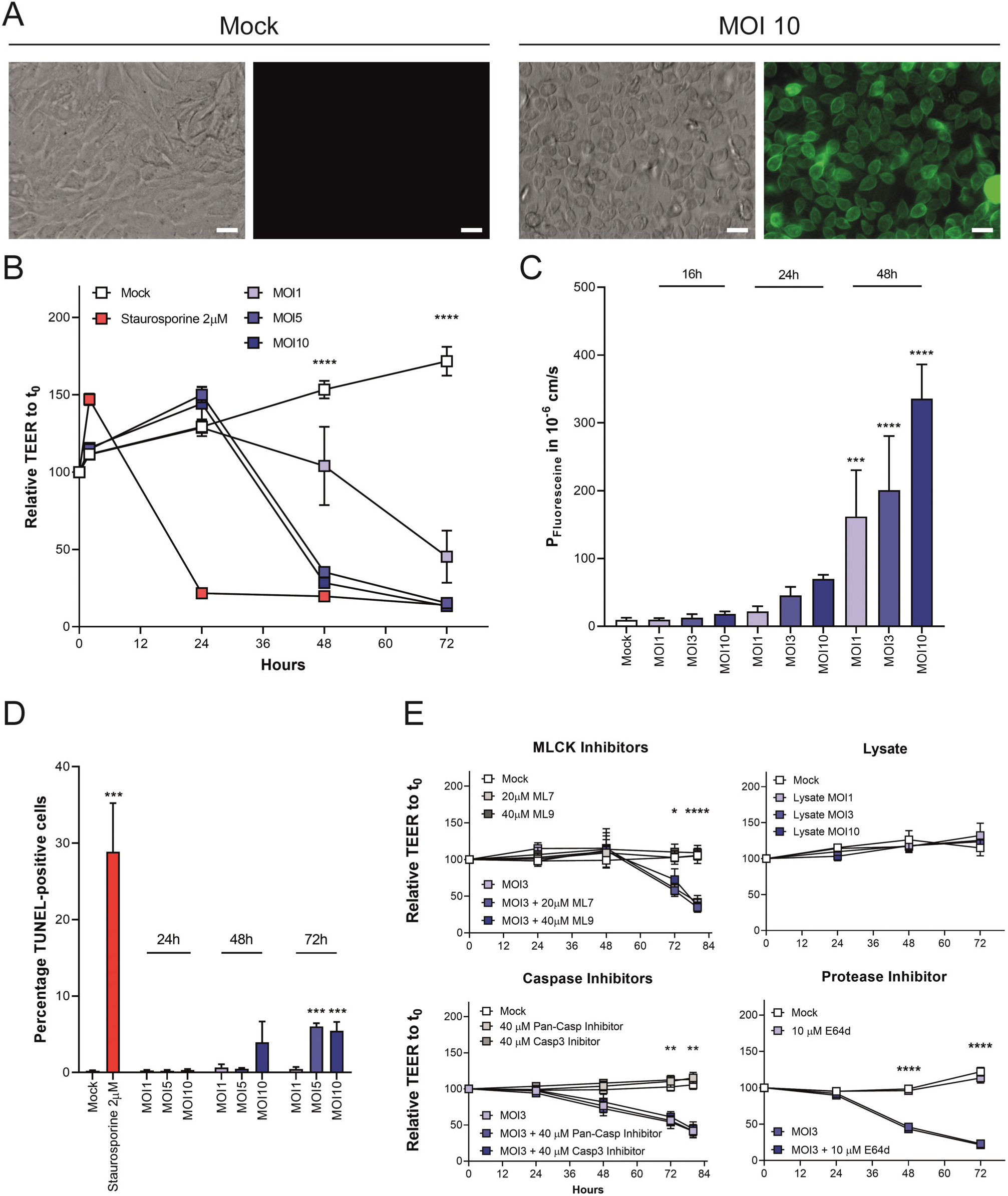
Barrier breakdown of human primary intestinal epithelium after infection with *G. duodenalis* trophozoites. Organoid-derived monolayers (ODMs) were infected with different parasite doses as indicated by different multiplicity of infection (MOI) and analyzed at the indicated time points after infection. (A) Representative light and fluorescent microscopic images of non-infected or MOI 10-infected ODMs with cell mask labelled *G. duodenalis* trophozoites (green) after 2 h of attachment. The amount of trophozoites in the MOI 10 condition reflects an epithelial cell layer completely covered with parasites. For comparison with other MOIs please refer to Supplementary Figure 2. Scale bars represent 20 μm. (B) Drop of transepithelial-electric resistance (TEER) after *G. duodenalis* infection indicated a disruption of barrier integrity in a dose- and time-dependent manner. (C) *G. duodenalis* infection induces leaking of apical medium to the basal compartment over time as indicated by Fluorescein (332 Da) translocation over the barrier. (D) Infection with *G. duodenalis* increased fraction of TUNEL-positive apoptotic cells in ODMs only at late time points post infection. (E) Factors previously proposed to potentially modulate barrier function in intestinal epithelia were tested for their capability to rescue the observed TEER decrease in ODMs after *G. duodenalis* infection. Neither Myosin light chain kinase (MLCK)-inhibitors (ML7 and ML9), nor caspase-inhibitors (pan-CASP and CASP3) or cysteine protease inhibitor (E64d) could rescue *G. duodenalis* induced TEER breakdown. Also, *G. duodenalis* lysate did not modulate TEER in ODMs. These results point towards alternative ways of modulation in ODMs. All quantitative experiments show mean (± SEM) of at least six individual transwell filters of at least two independent experiments. Statistical significance of treated samples in comparison to respective untreated mock control samples was determined using ANOVA with Dunnett’s correction for multiple testing. * p < 0.05, ** p < 0.01, *** p < 0.001 **** p < 0.0001

To study the effects of *G. duodenalis* on barrier function, ODMs were infected with trophozoites at a multiplicity of infection (MOI) ranging from 1 to 10 (Figure 2). At MOI of 10 parasites reached a near confluent density of attached trophozoites covering the ODM surface (Figure 2A, supplementary Figure 2). TEER decreased dose- and time-dependently after infection (Figure 2B). Loss of TEER, an integral measure of mainly paracellular barrier function, was reproducibly observed when infecting ODMs established from several individual donors (supplementary Figure 3). We also tested the effect of various *G. duodenalis* genotypes on TEER at MOI of 10 and found a similar pattern of TEER decrease after infection (supplementary Figure 4). Thus, we concluded that disruption of barrier function is unlikely dependent on a particular genetic background of host or parasite.

Paracellular barrier changes were also probed with the fluorescent tracer fluorescein (332 Da).^19^ Similar to the decrease in TEER, exclusion of fluorescein from the lower compartment was compromised in a MOI- and time-dependent manner (Figure 2C).

Using conventional epithelial cell line models, paracellular barrier and TEER breakdown due to *Giardia* has been linked to the action of parasite-released cysteine proteases and the activation of caspase-3 dependent apoptotic processes.^1, 20–23^ The caspase-3-dependent processes reported could be inhibited by myosin light-chain kinase (MLCK) inhibitors.^24^ Therefore, we tested whether TEER decrease in infected ODMs would also depend on these factors.

First, apoptosis of infected ODM was assessed by TUNEL staining 24, 48 and 72 h post infection. The frequency of apoptotic cells was not significantly different from uninfected control levels until 48 h post infection at MOIs of 1 and 5. Elevated numbers of TUNEL-positive cells were observable but only after 48 h post infection at MOI of 10 and later at 72 h post-infection at MOI of 5. Also, relative abundance rose to about 6% only which was just a fraction of the staurosporine-induced apoptotic cells in the respective control (Figure 2D). Thus, elevated numbers of apoptotic cells could only be observed after the TEER breakdown had already happened. In line with this finding, no measurable effect on *Giardia*-induced TEER breakdown was observed after addition of inhibitors of MLCK (ML7 or ML9) or caspase-3 (Pan-caspase and caspase-3 inhibitors) (Figure 2E, supplementary Figure 5).

To analyze the potential role of secreted cysteine proteinases that previously have been linked to barrier breakdown in cell line models^1, 21–23^ ODM were incubated with parasite lysates corresponding to the MOI range up to 10. Unexpectedly, this did not cause TEER decline (Figure 2D) despite high cysteine protease activity (supplementary Figure 6). The finding was corroborated in a reverse experiment where inhibition of cysteine proteases by E64d did not prevent TEER breakdown (Figure 2D). Notably, to exclude possible bias by decreasing viability of parasites during assays we could also show that trophozoite viability is not significantly altered by incubation with ODMs (supplementary Figure 7)

### *G. duodenalis* colonization of ODMs results in dramatic transcriptional responses in pathways for cell cycle control, cell-autonomous danger signal recording, ion transport and TJ structure

To discover early changes that may precede and cause TEER changes and barrier breakdown, we performed a transcriptome analysis at different time points post infection, but before TEER decline. ODMs were infected with a MOI of 5 and harvested after 1.5 and 24 h. Uninfected ODMs treated with similar media changes were lysed at the same time points and served as controls. Total RNA was reverse transcribed and cDNA libraries prepared for Illumina sequencing and transcript analyses were performed. Principle component analysis of relative transcript abundance values in samples analyzed 1.5 h after infection indicated no significant overall effect compared to non-infected controls (Figure 3A). In contrast, transcriptome profiles changed highly consistently in the five biological replicates with infection progressing to 24 h (Figure 3A). Next, pathway analysis of terms enriched in the annotation of differentially abundant mRNAs was performed in Enrichr using implemented gene-set libraries’ BioPlanet 2019 and GO terms (supplementary Figure 8). The analyses indicated signatures of effects on cell cycle control, chromatin organization, as well as on the epithelial cells’ innate immune response including NF-κB signaling (Figure 3B and supplementary data file 1). A reflection of this is seen in the annotations of the 40 most highly affected mRNAs that could also be verified via RT-qPCR (Figure 3C, supplementary Figure 9). Notably, transcripts of tumor necrosis factor-alpha (TNF-α) showed the largest relative increase due to infection (Figure 3C). We determined basolateral TNF-α protein secretion by ELISA and detected increasing amounts over time (supplementary Figure 10). In view of this, it was unexpected that claudin-2 mRNA was dramatically reduced (Figure 3C). However, the latter was consistent with an increase in TEER if ODMs were exposed to recombinant TNF-α basolaterally (supplemental Figure 11). This observation is also congruent with the recently described wound healing promoting effect of TNF-α in intestinal epithelia.^25^

**Figure 3:**
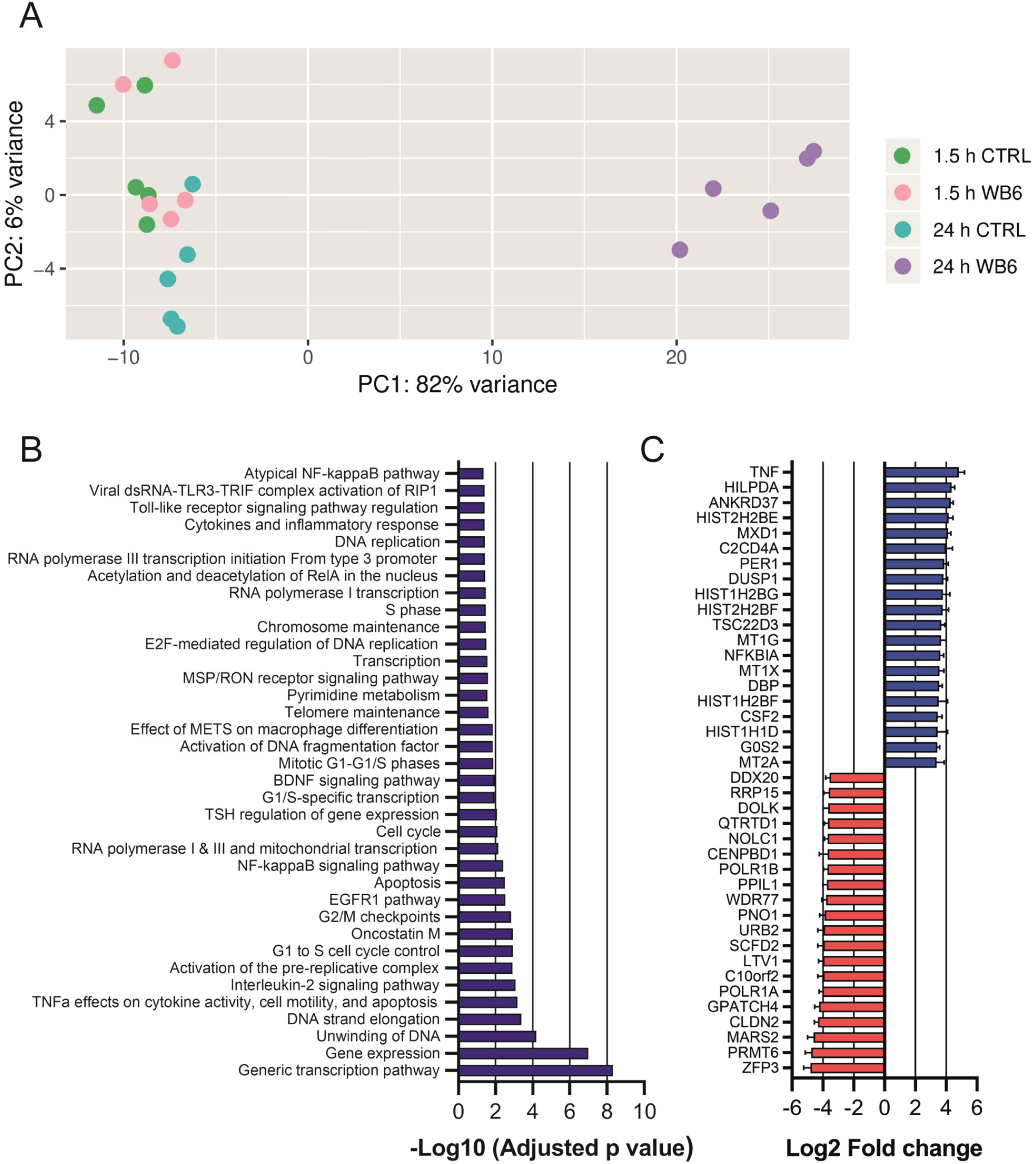
Altered ODM transcription during colonization with *G. duodenalis*. To determine possible factors preceding barrier breakdown in ODMs by *G. duodenalis* the transcriptome of ODMs was analysed by RNASeq in 5 independent replicates per condition at 1.5 and 24 h post infection (MOI of 5). At this time point and MOI barrier integrity was still intact. (A) Principle-component analysis showing distinctive differences in grouping of *G. duodenalis*-infected ODMs at 24 h post infection (24 h WB6) compared to mock infected control (1.5 h and 24 h CTRL) and *G. duodenalis*-infected ODMs at 1.5 h post infection (1.5 h WB6). (B) Significantly affected pathways, as deduced by gene ontology analysis, of ODMs infected with *G. duodenalis* at 24 h post infection compared to uninfected controls. Effects on cell cycle control, chromatin organization, and on innate immune response including NF-κB signaling are prominent (C) TOP 40-regulated genes of ODMs infected with *G. duodenalis* at 24 h post infection reflect this modulation. Data show mean (± SEM) of 5 independent biological replicas per condition.

Since TEER and paracellular transport will be affected by the relative contribution of claudins to TJs, we analyzed the transcriptome data for overall changes in mRNA abundances of claudins (Figure 4A, supplementary data file 2). ODM expressed claudin-1, −2, −3, −4, −7, −12, −15 and −18, which is a pattern consistent with that of the small intestine.^26^ Apart from claudin-2, infection for 24 h led with high consistency to roughly 2-fold changes in mRNA abundances for claudin-1 (reduced) and claudin-4, −7, −12 and −15 (all increased). Claudin-3 and −18 mRNA levels were not affected by infection. Overall, this indicated a strong effect of infection on claudins predictive of altered TJ channel (−2, −7, −12, −15) and barrier (−1) function. Transcripts of other components of the TJ complex include cytosolic adapters like TJP1 (ZO-1) proteins and structural transmembrane proteins such as occludin and were also investigated. At 24 h post infection relative TJP1 (ZO-1) mRNA abundance was only 1.4-fold reduced, while occludin mRNA was 2.7-fold more abundant than in controls (Figure 4A).

**Figure 4:**
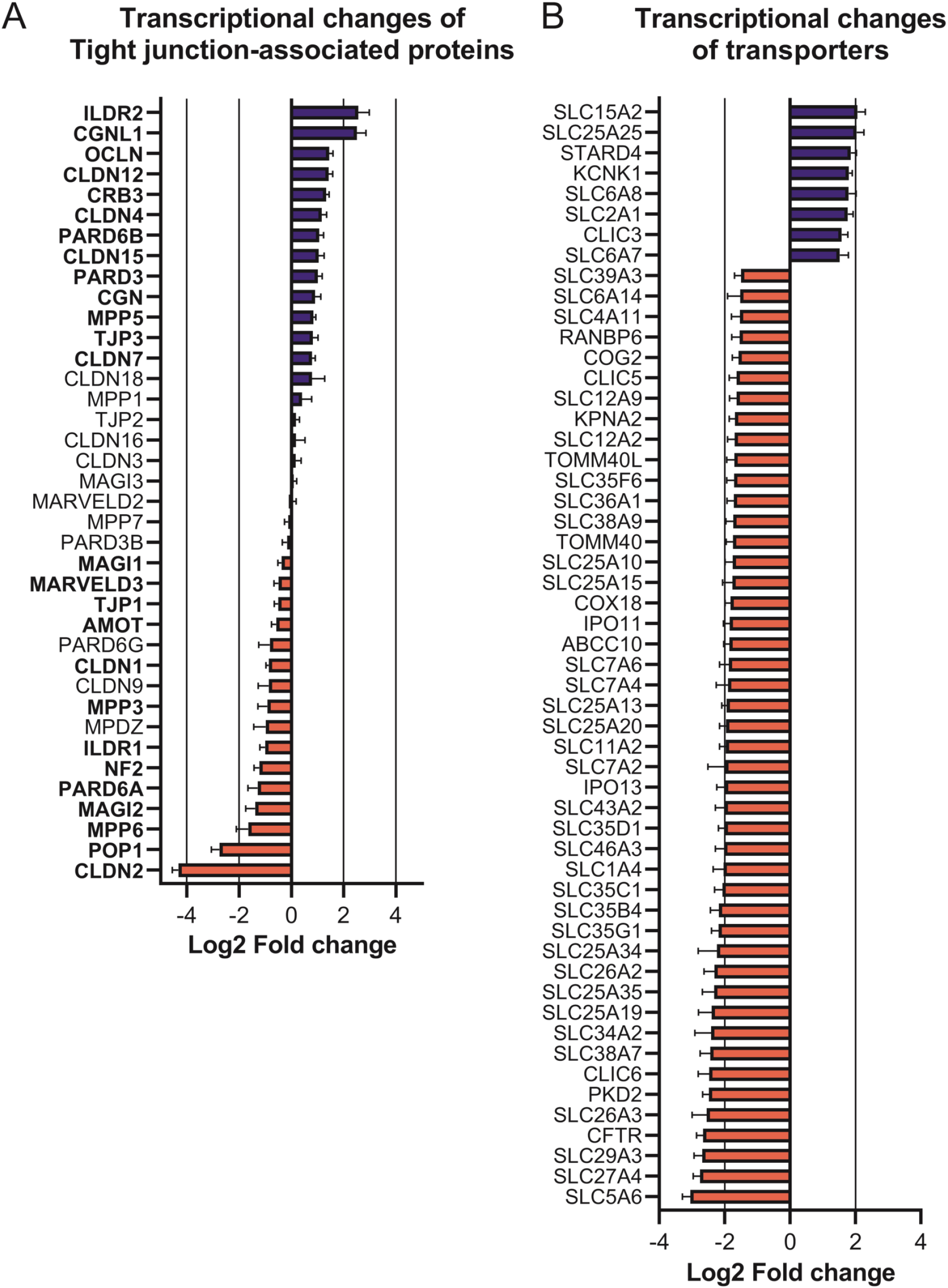
Alteration of gene expression of ion transporter and tight junction components in ODMs by *G. duodenalis*. Transcriptional changes of transporter and tight junction-associated genes 24 h post infection compared to mock control were analyzed from dataset shown in Figure 3 and supplementary data file 1. (A) Transcriptional changes of selected tight junction-associated genes are presented. Bold genes represent significant changes compared to mock infected controls. (B) Altered gene expression of solute carrier (SLC) family. Only significantly differently expressed transcripts of the SLC gene superfamily are shown (cut-off p<0.001 and >fold change of log2(1,5)). Data show mean (± SEM) of 5 independent biological replicas per condition.

The changes in claudin expression were consistent with the ensuing changes in TEER due to infection. However, in particular the decrease of claudin-2 mRNA in the presence of a TNF-α-NF-κB-activating cell intrinsic innate immune response was surprising as the contrary is observed in IBD.^27^ Reduction of claudin-2 transcript abundance was previously reported to be linked to reduced levels of the anion transporter SLCA26A3 (DRA).^28^ Thus, we mined the transcriptome data for changes in abundance of transporter mRNAs. Indeed, transporters were broadly and mostly negatively affected (Figure 4B). 24 h post infection SLCA26A3 (DRA) mRNA was ~6-fold reduced. Similarly, mRNAs encoding the basolateral SLC12A2 (NKCC1) and apical CFTR chloride channels were significantly less abundant.

This transcriptional response of ODM to infection suggests a disruption of ion transport processes preceding functional changes to TJ properties due to differential regulation of TJ protein expression. Eventually, the latter resulted in TEER and barrier breakdown and, ultimately, in disrupted ODMs. Thus, we aimed at seeking evidence in support of this hypothesis.

### *G. duodenalis* infection reduces SLC12A2 (NKCC1) and CFTR-dependent chloride secretion by human ODM and causes ultrastructural changes in the TJ network

Electrogenic chloride anion transport by SLC12A2 (NKCC1) was determined by stimulating secretion through addition of prostaglandin E2 (PGE_2_) to the basolateral compartment and theophylline to both, basolateral and apical compartment and determining the short circuit current (I_SC_) that can be inhibited with bumetanide, a selective inhibitor of SLC12A2 (NKCC1). Infection led to a reduction in the bumetanide-inhibitable fraction (Figure 5A). Importantly, this was observed in cultures already at 24 h post infection.

Triggering apical chloride secretion by forskolin and inhibiting the contribution of CFTR to this by addition of 5-nitro-2-(3-phenylpropylamino) benzoic acid (NPPB), revealed reduced CFTR activity in infected ODMs, in agreement with the transcriptomic data (Figure 5A).

**Figure 5:**
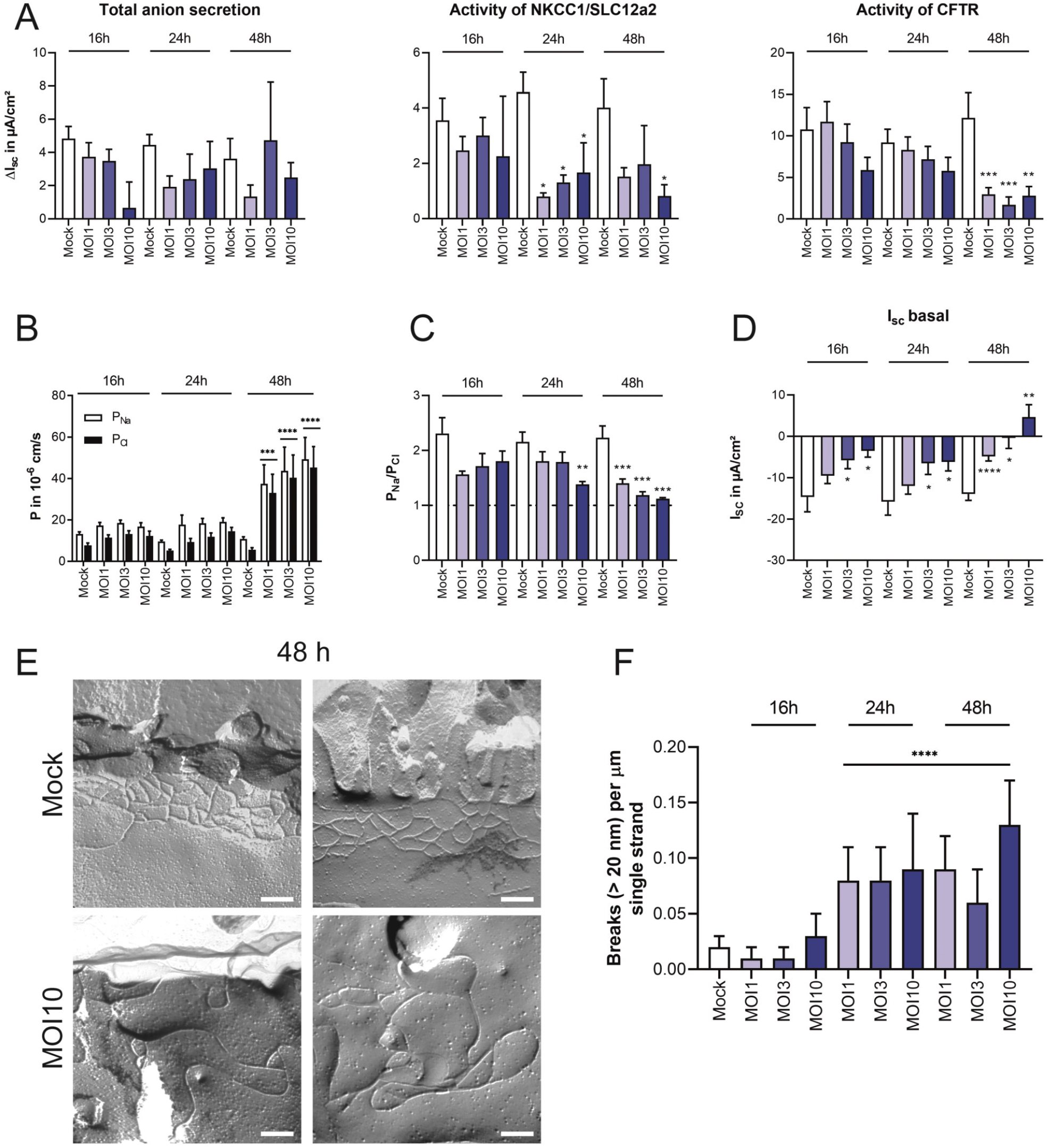
Alteration of electrophysiological properties of ODMs during *G. duodenalis* infection. Organoid-derived monolayers (ODMs) were infected with different parasite doses as indicated by different multiplicity of infection (MOI) and electrophysiological properties were analysed in an Ussing chamber set-up. (A) Assessment of the influence of *G. duodenalis* infection on total anion secretion and the activities of NKCC1/SLC12a2 and CFTR. All assays show a reduction of inhibitable fraction in comparison to controls supporting the results of the transcriptomic data, i.e., a loss of general and specific transporter activity. (B) Permeabilities for sodium (P_Na_) and chloride (P_Cl_) of infected ODMs and controls at indicated time points. While the control preserves low permeabilities, infected ODMs become leaky after 48h of infection. (C) Ratio of the permeabilities for sodium and chloride (P_Na_/P_Cl_). Controls preserve ratio ~2 over time while infected ODMs show dose- and time-dependent effects reaching ultimately a ratio of 1, i.e., a loss of ion-selectivity. (D) Basal short circuit current (I_sc_) determination showing changes in ion-transport of ODMs in a dose- and time-dependent manner. (E) Representative freeze-fracture electron microscopy (FFEM) images of infected (MOI 10) and non-infected (Mock) ODMs after 48 h. Infected ODMs show a loss of tight junctional organization only at late time points (48 h) and high MOIs. Scale bars represent 200 nm. (F) Quantification of breaks (> 20 nm) per μm of tight junctional strands using FFEM. Results show very early effect of infection leading to an accumulation of strand breaks already after 24 h - before breakdown of TEER. Statistical significance of treated samples against respective untreated mock control was determined using ANOVA with Dunnett’s correction for multiple testing. * p < 0.05, ** p < 0.01, *** p < 0.001, **** p < 0.0001. All electrophysiological experiments show means (± SEM) of at least five individual transwell filters of at least two independent experiments. FFEM data show mean (± SEM) of 20-43 TJ assessed per condition.

Paracellular ion permeability and charge selectivity were determined next by derivation from dilution potentials (Figure 5B, 5C). Uninfected ODM reproduced the cation-selective permeability of the small intestinal epithelium with a P_Na_/P_Cl_ ratio of roughly 2 (Figure 5C). During infection, absolute permeabilities for ions increased steadily and concomitantly the strong cation-selectivity of the ODMs were lost indicating an overall impairment of the TJ barrier (Figure 5D). These functional changes correspond directly to the reduced expression of claudins. Repressed expression of these major TJ components is often used as a predictive for reduced paracellular barrier function and loss of cation selectivity of the TJ network.

To assess ultrastructural consequences of the TJ modulation, we next performed freeze-fracture electron microscopy and analyzed changes in TJ ultrastructure. Infection led to an alteration of the ultrastructural TJ composition and a loosening of the TJ meshwork only at 48h post infection when paracellular barrier was lost already, which is in line with the electrophysiological results shown before (Figure 5E, supplementary Figure 12). It became also obvious that the localization of the TJ structures was shifted and only ~20% of the TJs were still located closely to the apically located microvilli. At earlier time points, the number of TJ strands, types and pattern, as well as meshwork depth and localization remained unaltered (supplementary Figure 12). A notable exception to this were changes in the increased number of single strand breaks that were observed already at 24h and persisted at 48h post infection (Figure 5F). This indicated early modulation of TJ organization.

### Infection alters TJ composition, localization and structural organization

To assess whether transcriptional changes in mRNA abundance and reduced barrier function also led to alteration of TJ complex at protein level, we performed immunofluorescence analyses of TJ components (Figure 6). For analysis the following set-up was chosen: Infections of ODMs were performed using MOI of 1, 3 and 10 and TEER was monitored over time. Cells were fixed when TEER values of the MOI 3 condition were reduced by ~50%. At this time point MOI 1 condition showed no TEER decrease and MOI 10 condition a complete TEER reduction. Therefore, this method allowed the analysis of alterations in the TJ complex in relation to TEER decline. Congruently with the reduced TEER values, we observed in the MOI 3 and 10 condition severe alterations in TJ proteins claudin-1 and −2, TJP1 (ZO-1) and occludin, which were reduced and delocalized from the junctional complex. In MOI 1 conditions, where TEER was not yet modulated, no alteration of TJP1 (ZO-1) and claudin-1 were detected, whereas claudin-2 was already less and, consistent with the transcriptomic data, occludin more abundant (Figure 6).

**Figure 6:**
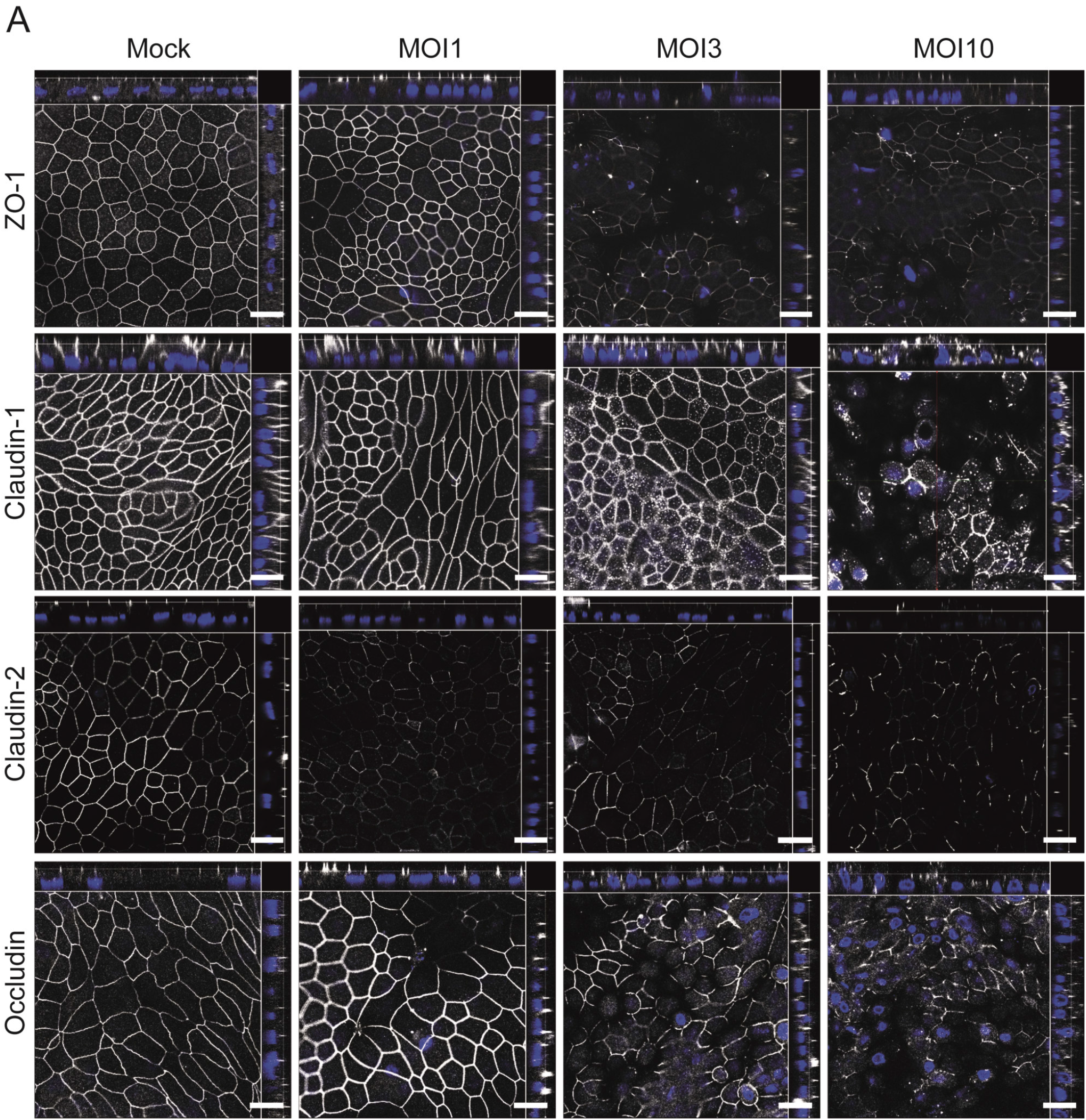
Modulation of the structure of tight junctional complexes in organoid-derived monolayers (ODMs) by *G. duodenalis*. ODMs were infected with different parasite doses as indicated by different multiplicity of infection (MOI) and monitored for structural components of the tight junction complex. *G. duodenalis* infected ODMs were fixed and analyzed by immunofluorescence microscopy when TEER values in the MOI3 sample dropped by 50% (images shown at 48 h). At this stage the MOI 1 fraction showed no TEER alteration, while the MOI 10 fraction showed complete TEER breakdown. Representative immunofluorescence images are shown as orthogonal stacks for tight junction proteins TJP1 (ZO-1), claudin-1, claudin-2 and occludin. Shown is MOI dependent modulation of tight junctional organization and delocalization of tight junctional proteins towards more basal regions. Scale bars represent 20 μm.

## Discussion

Here, an *in vitro* human duodenal organoid culture system was established to overcome limitations of classical cell lines in revealing the deleterious effects of colonization of functionally tight primary epithelial cell layers by *G. duodenalis* parasites. Parasites caused barrier breakdown independently of the previously reported pathways. Instead, barrier function loss was the culmination point of an early reduction of transporter transcript abundance and decrease of cellular ion transport activity that led over to para-cellular leakiness due to alteration of tight junction composition, localization and structure.

The specific pathophysiological correlates in the human host that lead to symptomatic outcome of giardiasis are not well understood. Most prominently, increased caspase-dependent apoptosis and tight junction disruption in intestinal epithelial cell lines has been linked to *Giardia*-induced increased epithelial permeability, ^1, 4, 20, 24, 29^ however, the sequence of events has not been studied in detail. In the seminal study that investigated changes in the mucosa of the small intestines of symptomatic individuals, a barrier dysfunction has been recognized and was correlated with alteration of the crypt villus length, decrease of epithelial resistance, increased permeability, altered ion and glucose transport, reduced claudin-1 expression and increased apoptosis.^8^ It was concluded that “leak flux, malabsorptive and secretory components” are key factors during chronic giardiasis. Our results are not only in good agreement with these *in vivo* findings but suggest a sequence of functionally linked events that lead to these changes.

In the ODM system we also found a parasite load-dependent leak flux indicated by collapse of epithelial electric resistance and increased transcellular translocation of fluorescein upon *Giardia* infection. Superficially, the data agree with findings that were generated using classical immortalized epithelial cell lines and animal models, and together point towards disruption of epithelial integrity by breakdown of the TJ complex as a major factor determining disease outcome.^1^ However, there are notable mechanistic differences between the findings using cell lines and the results reported here. In cell line models, the breakdown of TJ complexes was dependent on caspase-3 and myosin light-chain kinase (MLCK), and correlated with increased apoptosis.^1, 20, 24, 30^ Additionally, cysteine proteases were implicated in the loss of TJ integrity in this cell model.^1, 22, 23, 31^ In contrast, in the ODM system neither MLCK inhibition nor parasite lysates that retained high cysteine protease activity, led to any decay of TEER. We also detected increased rates of TUNEL-positive, apoptotic cells. However, these appeared only after collapse of TEER indicating that apoptosis may be a consequence of TJ breakdown and exposure to luminal content ^32^ rather than its cause.

The transcriptional responses of the ODM epithelia system to *G. duodenalis* infection showed good agreements with previous studies using Caco2 cell line-derived epithelia models in the aspects of epithelial cells’ innate immune reaction (NF-kappaB pathway and chemokine upregulation) and cell cycle arrest.^33, 34^ The colon cancer-derived Caco-2 cells responded comparatively faster and with a larger amplitude (c.f. supplementary data file 2), a feature possibly related to their origin (see below), the colon that has evolved facing a much higher microbial exposure. However, the comparative analysis revealed clear differences with respect to transcripts encoding TJ proteins, proteins involved in response to oxidative stress and those encoding solute and ion transporters (c.f. supplementary data file 2). These discrepancies are unlikely to be due to the use of different parasites representing the two human pathogenic genotypic groups used here and in the compared study^33^ since we observed consimilar patterns of TEER decrease after infection using different genotypes.

We report changes in TJ composition akin to the findings in symptomatic patients.^8^ TJ protein expression changes along the GI tract^26^ and cells - including immortalized cell lines - carry an imprint of gene expression related to their origin that includes claudins.^35^ This pertains also to the colon-derived Caco-2 cells that show colon adenocarcimona-linked alterations in claudin-expression.^36^ Loss of paracellular barrier function correlating with a preceding decrease in claudin-1 and −2 mRNA levels and subsequent protein downregulation, as reported here using ODM, was not observed in Caco2 cells which as mentioned before is likely due to their ‘memory’ of their origin. In agreement with our observations, patients suffering from symptomatic giardiasis had claudin-1 downregulated and claudin-2 was undetectable.^8^

Notably, claudin-2 was decreased while TNF-α mRNA and protein levels were increased after infection which is in contrast to findings in inflammatory diseases such as IBD^27^ where additional pro-inflammatory cytokines are present. Using recombinant TNF-α and adding it to the system we also did not observe a decrease in TEER in uninfected epithelia but rather the opposite. This is in line with a recent study that highlighted the protective role of TNF-α receptor signaling in epithelial cells by direct effect of TNF-α on intestinal stem cells.^25^ It is therefore worth speculating that autocrine TNF-α produced after infection in intestinal epithelia might also play a protective role during giardiasis.

A number of studies and, in particular, *in vivo* analyses have indicated defects in ion homeostasis in epithelia after *G. duodenalis* infection.^37–42^ One of these studies, for example, reported a decrease in CFTR function in cells exposed to parasites or concentrated parasite secreted/released factors.^37^ We show that *G. duodenalis* infection alters ion transporter expression and function in epithelial cells before TJ modulation. Altered transporter expression has been shown to regulate changes in TJ composition and function.^43^ We further discovered that altered cellular ion transport precedes disruption of the paracellular barrier via ultrastructural and TJ protein expression changes and localization shifts, which ultimately causes cell death and epithelial disintegration. These findings with primary human duodenum-derived ODMs are hard to reconcile with the proposed viewpoint which states the reverse order of these events as the cause of symptomatic giardiasis.^1^ Methodological differences of the various models may explain some of the opposing findings. The recapitulation of *in vivo* observations in symptomatic patients, however, argues in favour of the ODM model. Moreover, CRISPR/Cas methodology^44^ adds reverse genetic tractability to the ODM model on different donor genetic backgrounds, opening the avenues to elucidate the causal chain of the underlying molecular mechanism(s) that link to symptomatic giardiasis, even on different host genetic backgrounds.

## Acknowledgments

The authors thank Gudrun Kliem, Antonia Müller, and Jasmin Gerkrath from Robert Koch-Institute and In-Fah M. Lee from the Institute of Clinical Physiology (Charité) for excellent technical assistance. We thank Scott C Dawson for reading and commenting the manuscript and for valuable discussions.

**Supplementary material**

Supplementary methods and figures

Supplementary material data file 1

Supplementary material data file 2

## Footnotes

## Data availability statement

All data are available in the original manuscript and online supplementary material.

